# Improved peak-calling with MACS2

**DOI:** 10.1101/496521

**Authors:** John M. Gaspar

## Abstract

The computational analyses of genome-enrichment assays, such as ChIP-seq and ATAC-seq, are typically concluded with a peak-calling program that identifies genomic regions that are significantly enriched. The most popular peak-caller, MACS2, assumes that the input alignment files are for single-end sequence reads by default, yet those with paired-end Illumina sequence data frequently use this default setting. This leads to erroneous coverage values and suboptimal peak identification. However, using the correct paired-end mode can introduce another set of artifacts. After thoroughly reviewing the MACS2 source code, we have modified it to limit these and other problems. Our updated version is freely available (https://github.com/jsh58/MACS).

---

Genome-enrichment assays, such as ChIP-seq, MeDIP-seq, ATAC-seq, and DNase-seq, involve the selection of short DNA fragments, either by immunoprecipitation using an antibody targeting a protein of interest, or by an enzymatic reaction that preferentially occurs in open regions of chromatin. Once obtained, the fragments are subjected to high-throughput sequencing, typically by an Illumina machine. One characteristic of Illumina sequencing technology is its ability to generate paired sequence reads, one from each end of a DNA fragment. This allows one to align the paired-end reads to a reference sequence and infer the original fragment’s extent without needing to sequence its entirety.

With genome-enrichment assays, the sequencing and read alignment steps are usually followed by peak-calling, in which genomic regions that are demonstrated to be significantly enriched are identified. The most popular peak-calling program is MACS2 [1, 2], with over 3,000 citations as of Oct. 2018. One of its attributes, inherited from its predecessor (MACS), is modeling DNA fragment lengths from single-end sequencing in order to extend the reads to represent the full fragments. Although this modeling is superfluous with paired-end sequencing, it remains the default analysis option.

Therefore, it has been surprising to note the large number of studies that have included genome-enrichment assays and paired-end sequencing but analysis by MACS2 in the default, single-end mode. A brief survey of papers from May-June 2018 revealed more than a half dozen such publications [3-8]. One paper [9] cited a ChIP-seq analysis pipeline [10] that explicitly recommends the single-end analysis mode of MACS2 (‘-f BAM’), even when the sequencing is paired-end.

We have carefully reviewed the MACS2 source code to better understand how the program interprets the input alignment file(s). When paired-end datasets are analyzed in single-end mode, MACS2 eliminates the second read of each pair (the “R2” read) and then treats the remaining “R1” reads as if they were single-ended. As described above, it models the fragment lengths from the “single-end” R1 reads and then extends the read lengths to the average value from the model. One can skip this modeling via a command-line option (‘--nomodel’), in which case the reads will instead be extended to a given length (default 200bp; see Supplementary Fig. 1).

Therefore, analyzing paired-end datasets with MACS2 in its default, single-end analysis mode fails to utilize all of the information contained in the datasets. At least half of the sequence data is disregarded automatically; all paired reads that are aligned in a “not properly paired” configuration are ignored as well.

Furthermore, except in the unlikely case that all of the assay’s selected DNA fragments are of uniform size and match the extension length, the fragment coverage values calculated by MACS2 would be erroneous. These coverage values are the primary determinant of peak identification by MACS2. Therefore, we hypothesized that the incorrect analysis mode would also decrease the quality of the peaks, especially in cases where a substantial number of the fragments’ lengths differed from the extension length.

We tested this hypothesis on a recently published dataset. The paper from Corces *et al*. [11] explicates a modification of the ATAC-seq [12] laboratory protocol, termed “Omni-ATAC.” The principal benefit of this technique is the limitation of mitochondrial-derived sequence reads. Such reads are of minimal interest to those conducting ATAC-seq experiments; they are routinely discarded in the computational analysis and thus represent a waste of sequencing resources.

The computational analysis performed by Corces *et al*. included peak-calling by MACS2 in the default, single-end mode, despite the sequencing being paired-end. Thus, the R2 reads were ignored, and, since the ‘--nomodel’ option of MACS2 was specified, the R1 reads were extended to 200bp. This would seem to be a suboptimal choice for these datasets, since the DNA fragment lengths followed a characteristic distribution [12], with just 21.7% of fragments having lengths within 10% of 200bp (Fig. 1A, Supplementary Fig. 2). Therefore, the fragment coverage values calculated by MACS2 were frequently in error (Fig. 1B, Supplementary Fig. 3). More than 40% of all genomic positions at which the true coverage value was at least five had calculated values that were incorrect by at least 50%.

**Figure 1.**
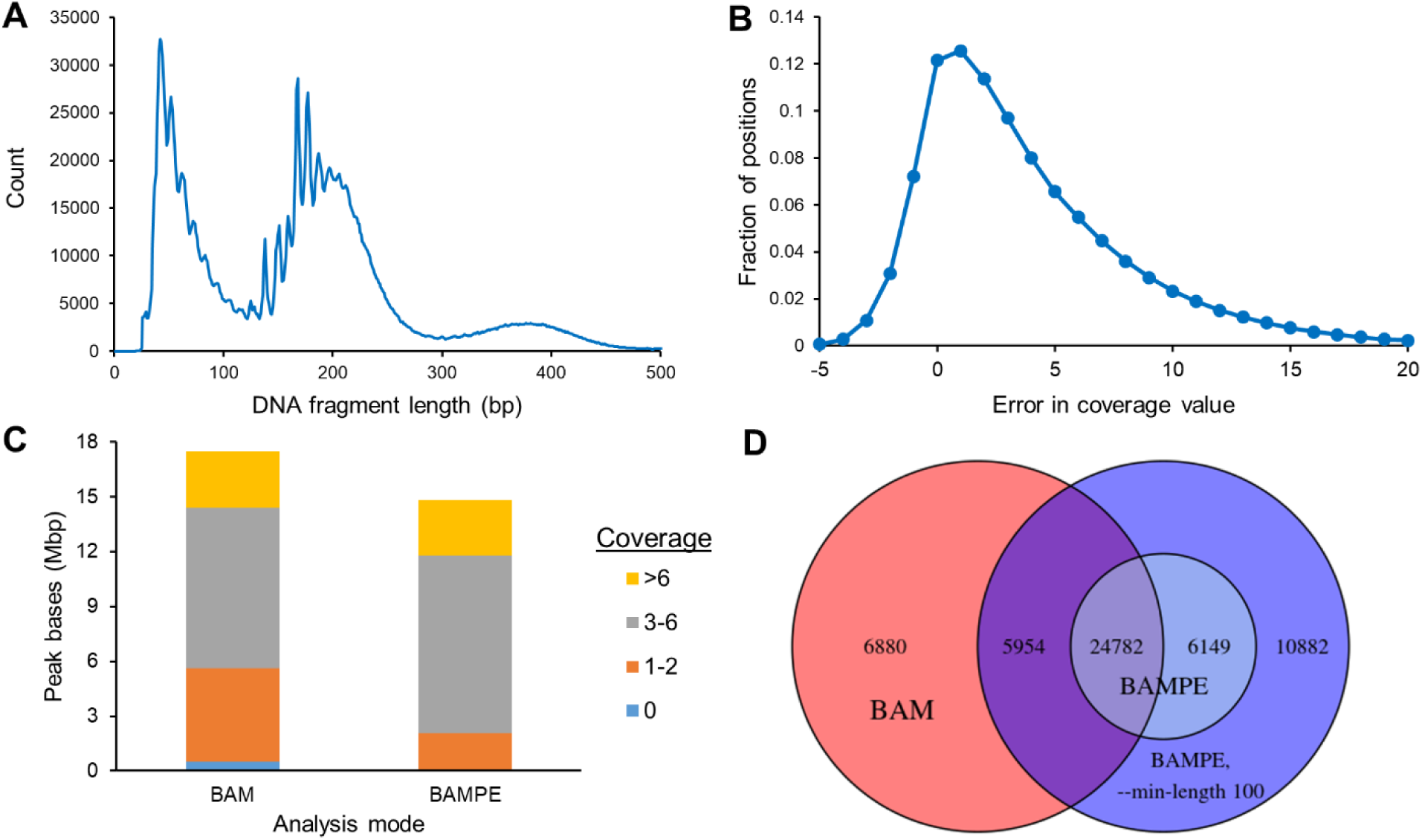
Results for sample SRR5427884. **A:** The distribution of fragment lengths followed a characteristic shape that is typically seen with ATAC-seq datasets. **B:** For all genomic positions at which the true fragment coverage (pileup) was at least five, MACS2 calculated the coverage accurately only 12% of the time. This was because, in single-end mode, MACS2 extends all R1 reads to 200bp, regardless of the true fragment length. **C:** Fragment coverages of peaks in single-end analysis mode (‘BAM’) versus paired-end mode (‘BAMPE’) of MACS2. **D:** A comparison of peaks called in single-end mode (‘BAM’), paired-end mode (‘BAMPE’), and paired-end mode using the modified MACS2 with a minimum peak length of 100bp (‘BAMPE --min-length 100’). Each peak set was compared against the union of all three peak sets.

The incorrect coverage values had a direct effect on the peaks called. For example, of the 17.5Mbp peak bases, 32.2% had true fragment coverages of less than three (Fig. 1C). The low-coverage peak regions in single-end mode occurred both in the vicinity of true peaks (Supplementary Fig. 1) as well as sites where there was no apparent enrichment (Supplementary Fig. 4).

In its paired-end analysis mode (‘-f BAMPE’), MACS2 interprets the full extent of the sequenced DNA fragments correctly (Supplementary Fig. 1). Although this resulted in fewer overall peaks (Supplementary Table), there were also reductions in the peak bases at which there were low fragment coverages (Fig. 1C). Nearly 1Mbp of additional peaks at coverages greater than two were called.

Although using the paired-end mode of MACS2 reduces apparent false positive peaks, some false negatives are actually more likely to occur. The reason is that, in paired-end mode, the minimum length of called peaks is set to the average fragment length calculated by the program. Therefore, any region that achieves statistical significance but fails to meet this length requirement is automatically discarded (Supplementary Fig. 5). We have modified MACS2 to accept two additional command-line arguments that allow the user to determine the thresholds for peak-calling (see Methods for details). We note that reducing the minimum length parameter to 100bp led to increased overlap between single-end and paired-end analysis modes (Fig. 1D, Supplementary Fig. 6).

Our purpose here is not to disparage the value of the Omni-ATAC protocol espoused by Corces *et al*.; indeed, it is apparent from these datasets that the protocol reduces mitochondrial contamination and is therefore beneficial. Rather, we have used these datasets to illustrate what appears to be the common practice of using MACS2 in an analysis mode that is inappropriate to the datasets, leading to suboptimal peak identification. This can confound further biological interpretation (e.g. transcription factor motif enrichment).

Those who have been dissatisfied with the results produced by MACS2 in paired-end mode may find that our introduction of two command-line parameters yields improved outcomes. We have made additional modifications to the MACS2 code to address other issues that we have discovered in our extensive review of it (see Methods, Supplementary Fig. 7). The source code of our version is freely available on GitHub (https://github.com/jsh58/MACS).

## Methods

Raw fastq files were downloaded from the NCBI SRA (accession SRP103230). The Supplementary Table contains details of the analyzed datasets.

Adapters were removed from the paired-end reads with NGmerge v0.2 [13], using default parameters except for the following: ‘-a -e20 -u41’. Reads were aligned to the human genome (hg19) by Bowtie2 v2.3.2 [14] with default parameters except for ‘-X 2000’. The following sets of alignments were removed: those not properly paired, putative PCR duplicates (properly paired fragments that shared both leftmost and rightmost genomic coordinates), alignments to chrM and chrY, and those with MAPQ less than 30. The alignments were converted to BAM with SAMtools v1.5 [15].

Peaks were identified with MACS2 v2.1.1.20160309 [1, 2] with the following arguments: ‘--keep-dup all --nolambda --bdg’. For analyses matching Corces *et al*. [11], with the default input format (‘-f AUTO’, corresponding to ‘-f BAM’), ‘--nomodel’ was specified. For paired-end analyses, ‘-f BAMPE’ was set. With the modified version of MACS2 (v2.1.2_dev), the ‘--min-length’ parameter was set to 100bp and the ‘--max-gap’ parameter was kept at the value used by the original version of MACS2 (i.e., the average fragment length for the sample).

BEDTools v2.26.0 [16] was used to combine peak sets and pileup coverage counts and to compute intersections and differences of peak sets. For each sample, the three peak sets (‘BAM’, ‘BAMPE’, and ‘BAMPE --min-length 100’) were compared against the union peak set created by merging the three peak sets (via ‘bedtools merge’). A union peak that overlapped peaks in multiple sets was counted as shared, regardless of the extent of overlap.

Genome visualizations were performed with IGV v2.4.9 [17].

The computations in this paper were run on the Odyssey cluster supported by the FAS Division of Science, Research Computing Group at Harvard University.

### MACS2 modifications

The following modifications were made to MACS2 (v2.1.1.20160309):

1. Two command-line arguments were added to the ‘macs2 callpeak’ function:
  - --min-length’: this parameter sets the minimum length required for a peak to be called (default: 100bp)
  - ‘--max-gap’: this parameter sets the maximum gap between significant sites to cluster them into the same peak (default: 50bp)
2. The miscalculation of average fragment length in the paired-end mode of ‘callpeak’ was corrected.
3. The miscalculation of *p*-scores (and thus *q*-scores also) due to the failure to resolve hash collisions was corrected for both ‘callpeak’ and ‘bdgcmp’.
4. The miscalculation of *p*-scores for large lambda (control) values was reduced. The ‘callpeak’ code was redirected to use ‘bdgcmp’ functions, which employ double floating-point precision (rather than single precision).

## Data and Software Availability

Raw fastq files were downloaded from the NCBI SRA (accession SRP103230).

The modified MACS2 (v2.1.2_dev) is freely available on GitHub (https://github.com/jsh58/MACS).

## Supplementary Information

**Supplementary Table:**
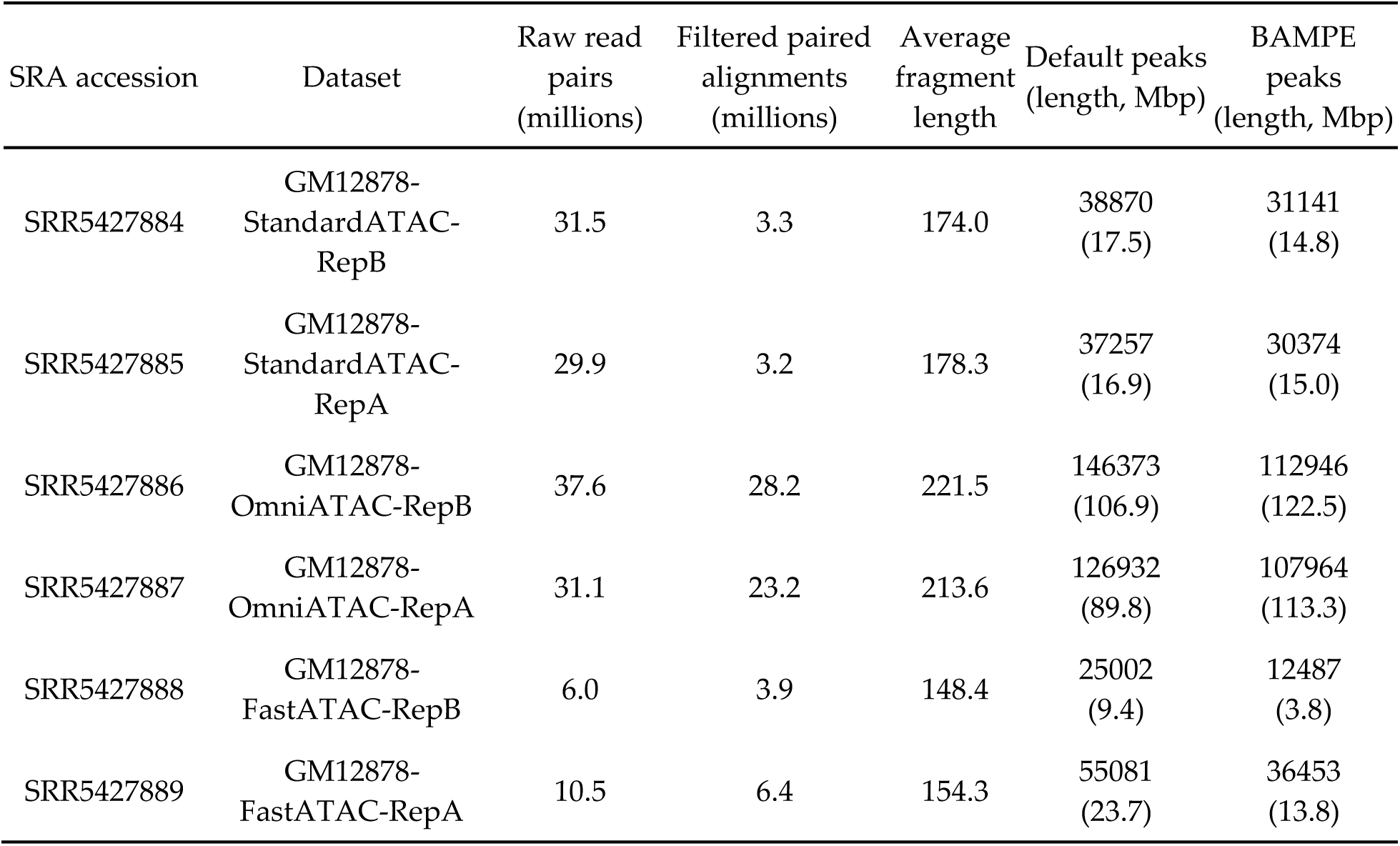
Details of datasets analyzed

| SRA accession | Dataset | Raw read pairs<br>(millions) | Filtered paired alignments<br>(millions) | Average fragment length | Default peaks<br>(length, Mbp) | BAMPE peaks<br>(length, Mbp) |
| --- | --- | --- | --- | --- | --- | --- |
| SRR5427884 | GM12878-StandardATAC-RepB | 31.5 | 3.3 | 174.0 | 38870<br>(17.5) | 31141<br>(14.8) |
| SRR5427885 | GM12878-StandardATAC-RepA | 29.9 | 3.2 | 178.3 | 37257<br>(16.9) | 30374<br>(15.0) |
| SRR5427886 | GM12878-OmniATAC-RepB | 37.6 | 28.2 | 221.5 | 146373<br>(106.9) | 112946<br>(122.5) |
| SRR5427887 | GM12878-OmniATAC-RepA | 31.1 | 23.2 | 213.6 | 126932<br>(89.8) | 107964<br>(113.3) |
| SRR5427888 | GM12878-FastATAC-RepB | 6.0 | 3.9 | 148.4 | 25002<br>(9.4) | 12487<br>(3.8) |
| SRR5427889 | GM12878-FastATAC-RepA | 10.5 | 6.4 | 154.3 | 55081<br>(23.7) | 36453<br>(13.8) |

**Supplementary Figure 1.**
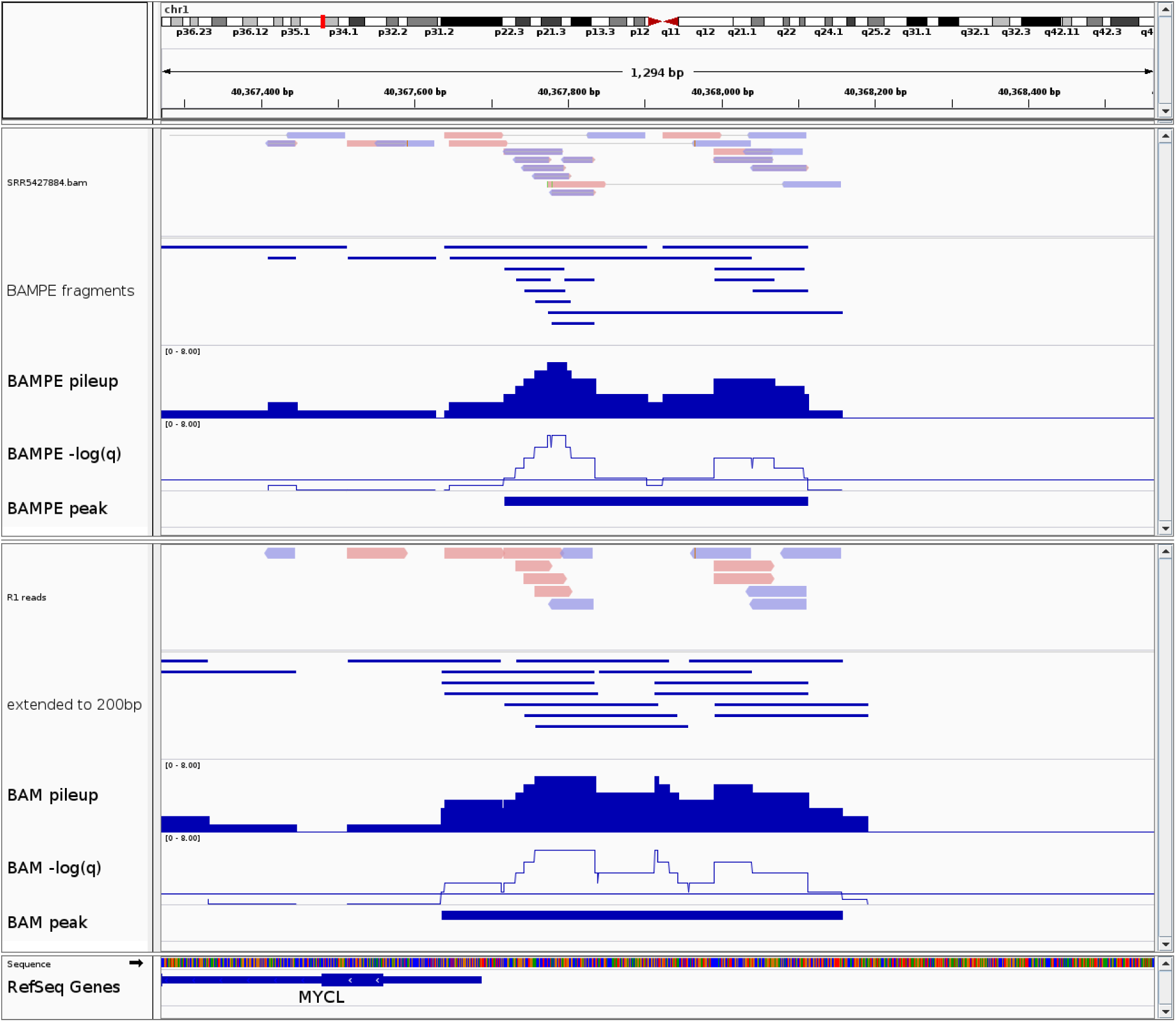
Sample peaks called in the different analysis modes of MACS2. For sample SRR5427884, the BAM file (top panel) showed a set of filtered, paired alignments near the transcription start site of *MYCL* on chromosome 1. In paired-end mode (‘-f BAMPE’, upper panels), MACS2 inferred the treatment pileup values from the full fragment lengths. All positions at which *q* < 0.05 (-log_10_(*q*) > 1.301) were considered significantly enriched, and a 396bp peak resulted. In the default, single-end mode (‘-f BAM’, lower panels), the R2 reads were eliminated and the R1 reads were extended to 200bp to represent the DNA fragments. This resulted in higher pileup values at most positions and a larger 523bp peak. Note that more than 10% of this peak included positions at which the true coverage was 0 or 1.

**Supplementary Figure 2.**
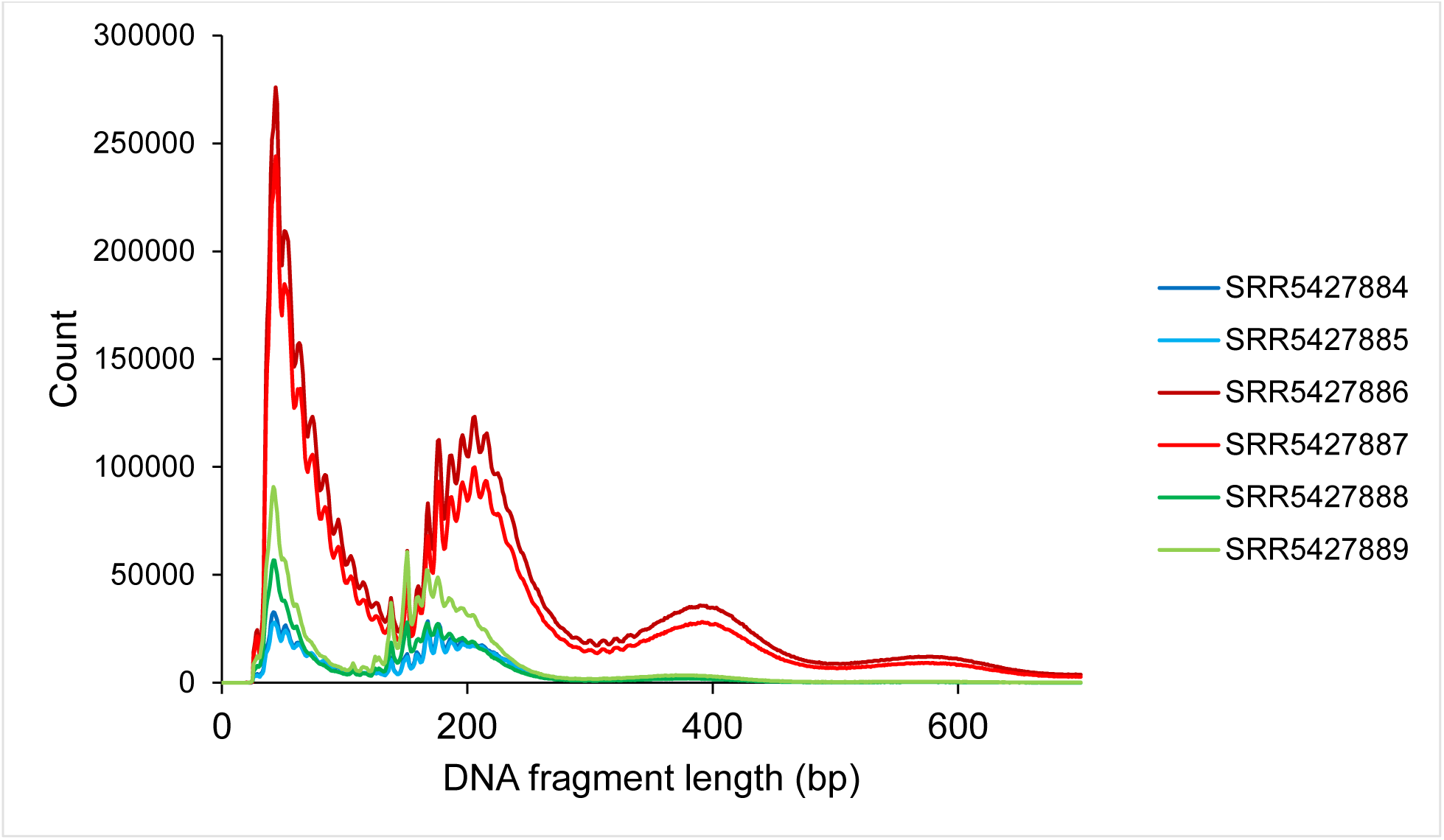
DNA fragment lengths of all six samples. After filtering the aligned reads, each sample showed the characteristic distribution of fragment lengths that is typical in ATAC-seq experiments.

**Supplementary Figure 3.**
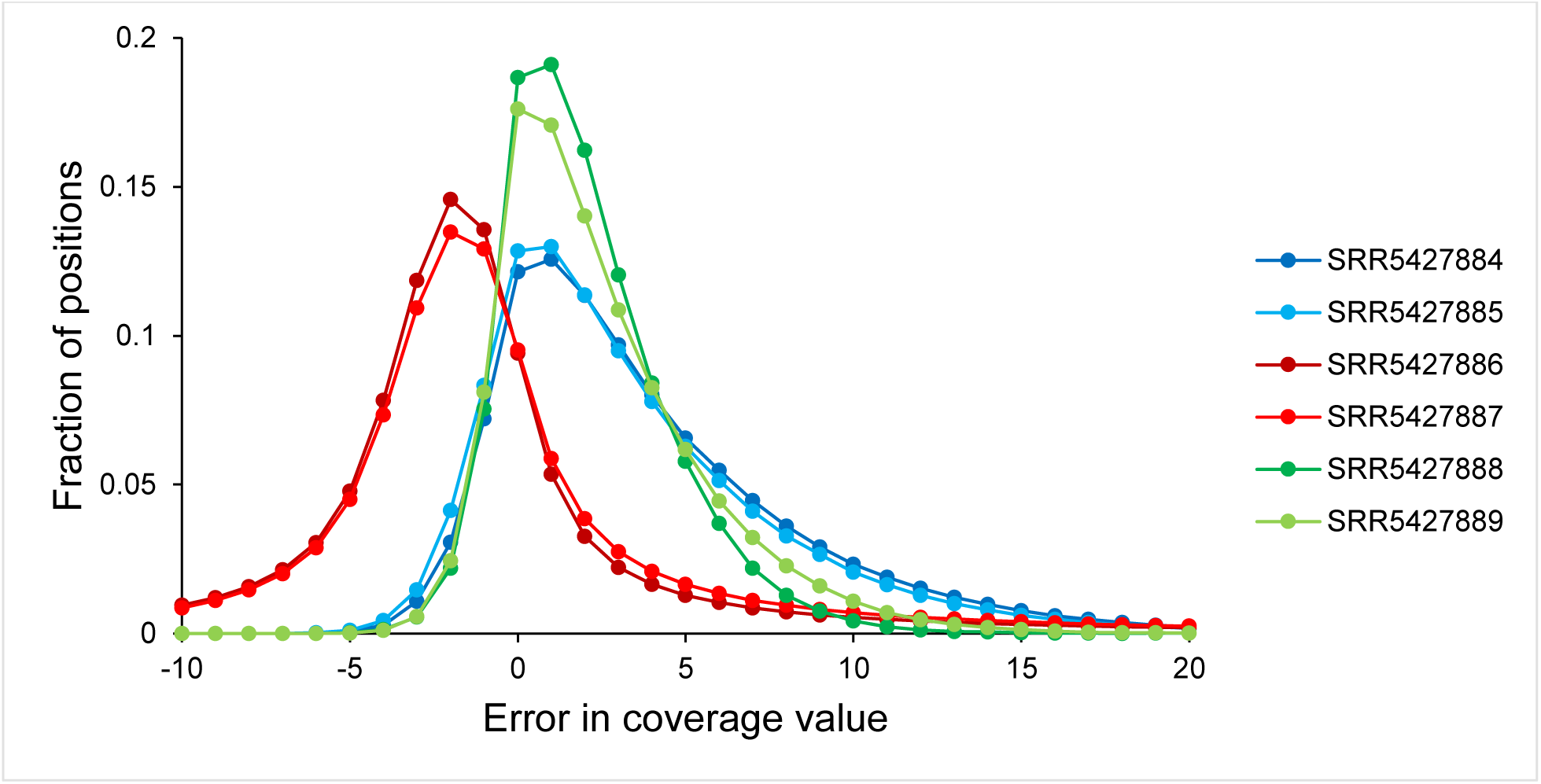
Errors in coverage values in single-end mode. Because of the extension of R1 reads to 200bp in single-end mode, the fragment coverage values calculated by MACS2 were frequently in error. Note that, for samples SRR5427886 and SRR5427887, there were more negative errors because the average fragment lengths of these samples were greater than 200bp (Supplementary Table).

**Supplementary Figure 4.**
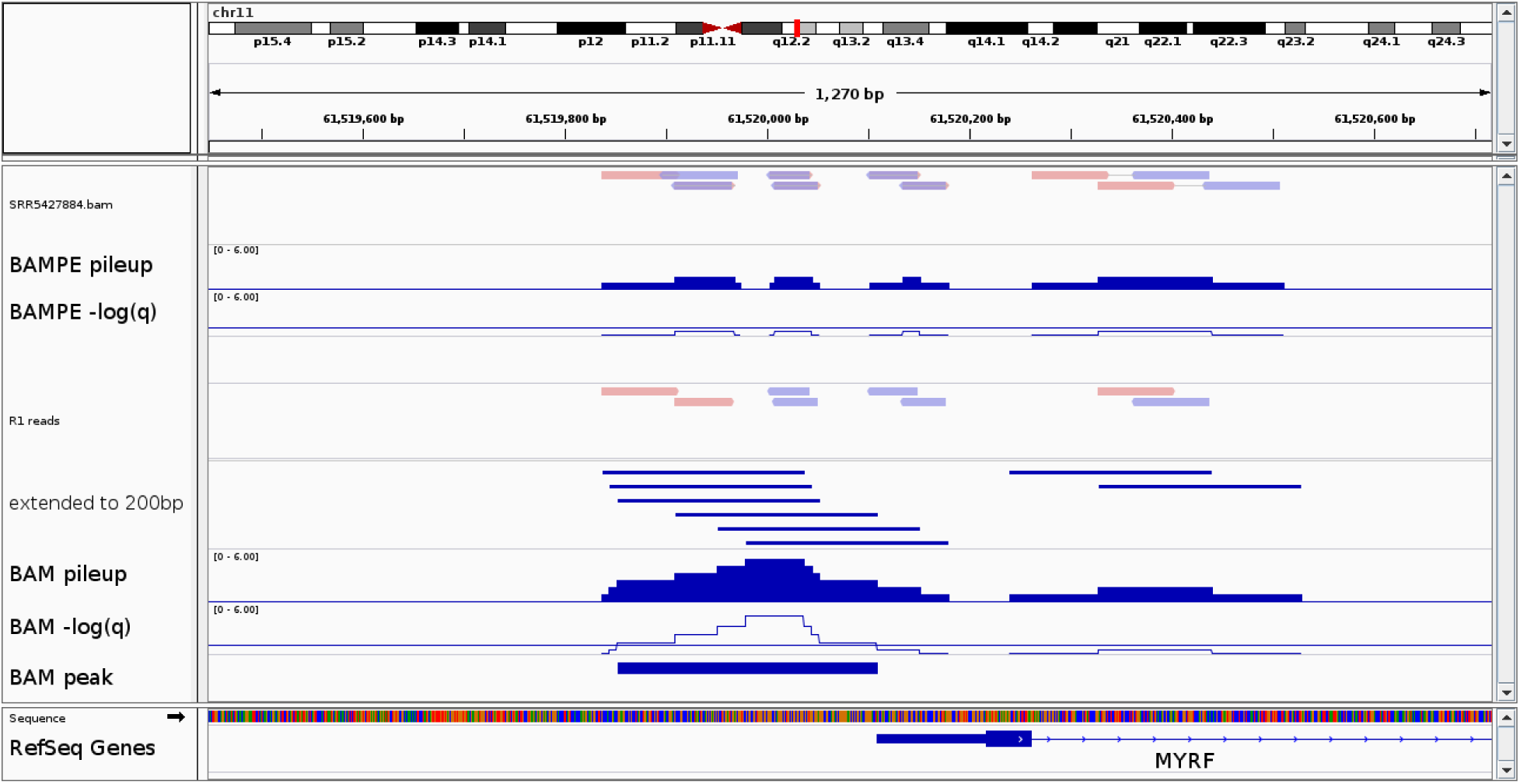
A false positive peak called by MACS2 in single-end mode. For sample SRR5427884, in paired-end mode (BAMPE, upper panels), the mostly short fragments were interpreted correctly, resulting in low pileup values and no *q*-values that reached statistical significance at any position in this region near the transcription start site of *MYRF*. A peak would have been called only if the statistical threshold (*q* < 0.05) were relaxed. In single-end mode (BAM, lower panels), the R1 reads were extended to 200bp, resulting in higher pileups that yielded significant *q*-values and a peak of 257bp.

**Supplementary Figure 5.**
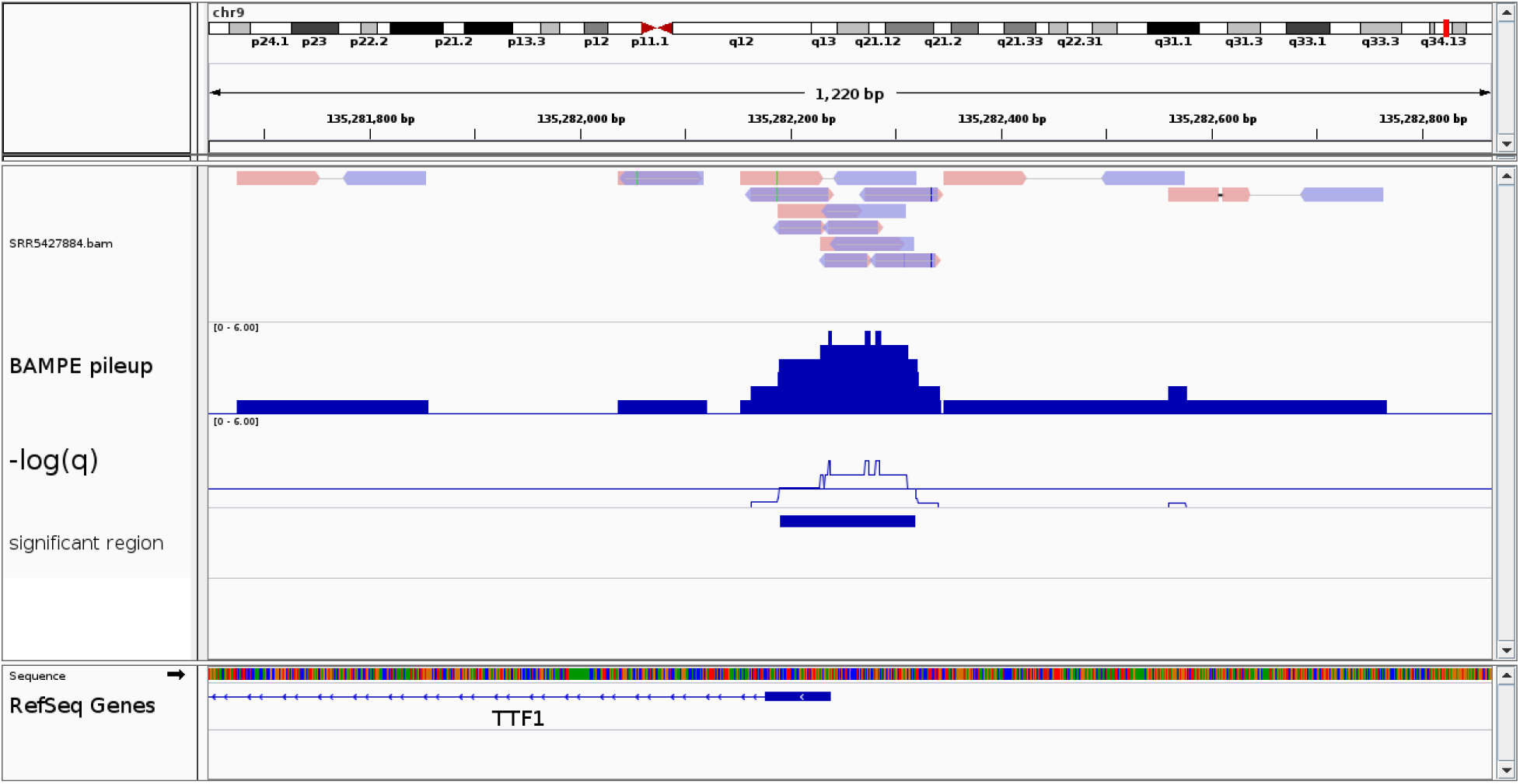
A false negative peak in paired-end mode. For sample SRR5427884, a 129bp region overlapping the transcription start site of *TTF1* achieved statistical significance. However, this region was discarded by MACS2 because the minimum peak length for this sample was 174bp (the calculated average fragment length). With our modified version of MACS2, the minimum peak length is a parameter (default 100bp), and a peak would have been called here if that parameter were less than or equal to 129bp.

**Supplementary Figure 6:**
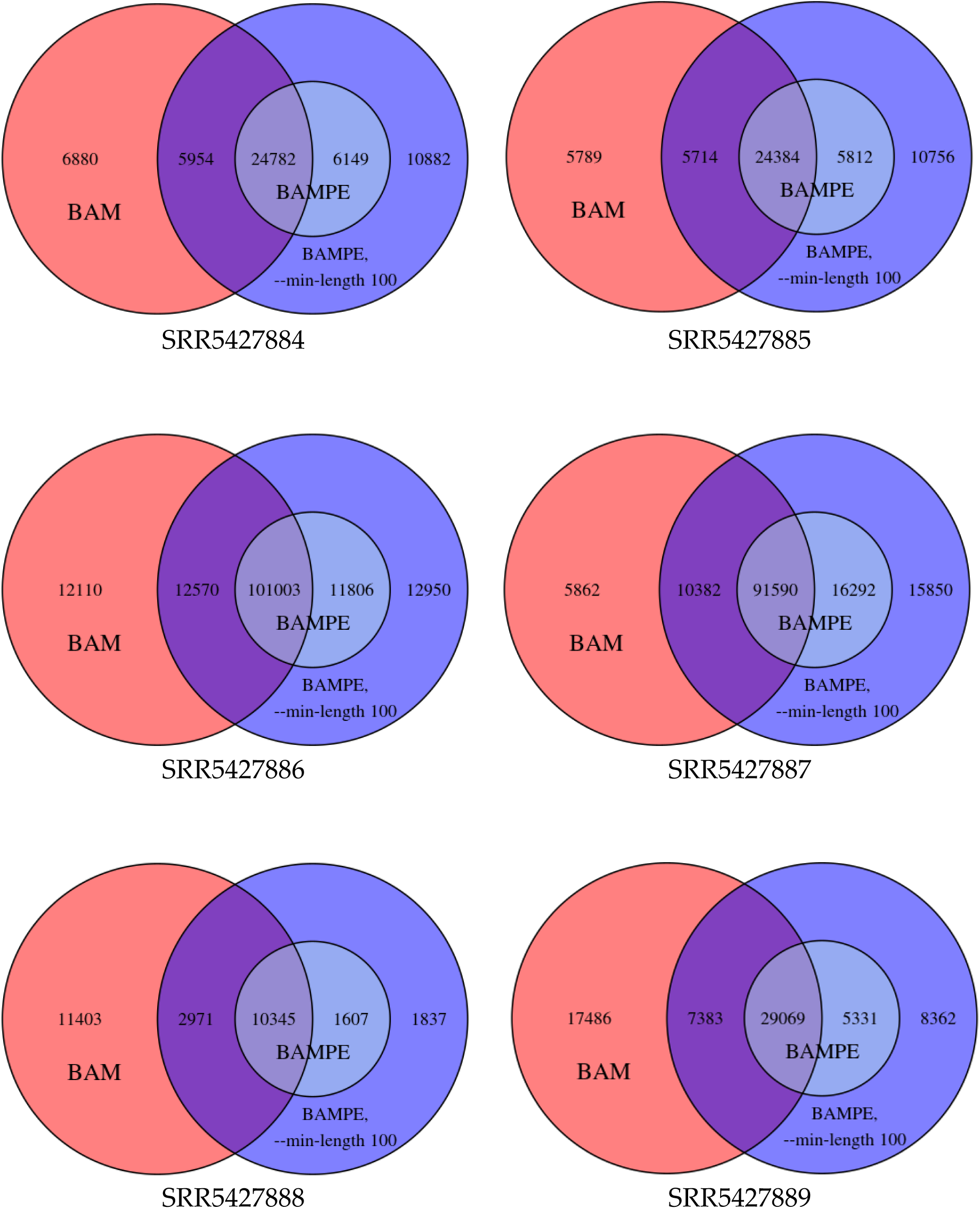
Peak overlaps in different analysis modes.

**Supplementary Figure 7.**
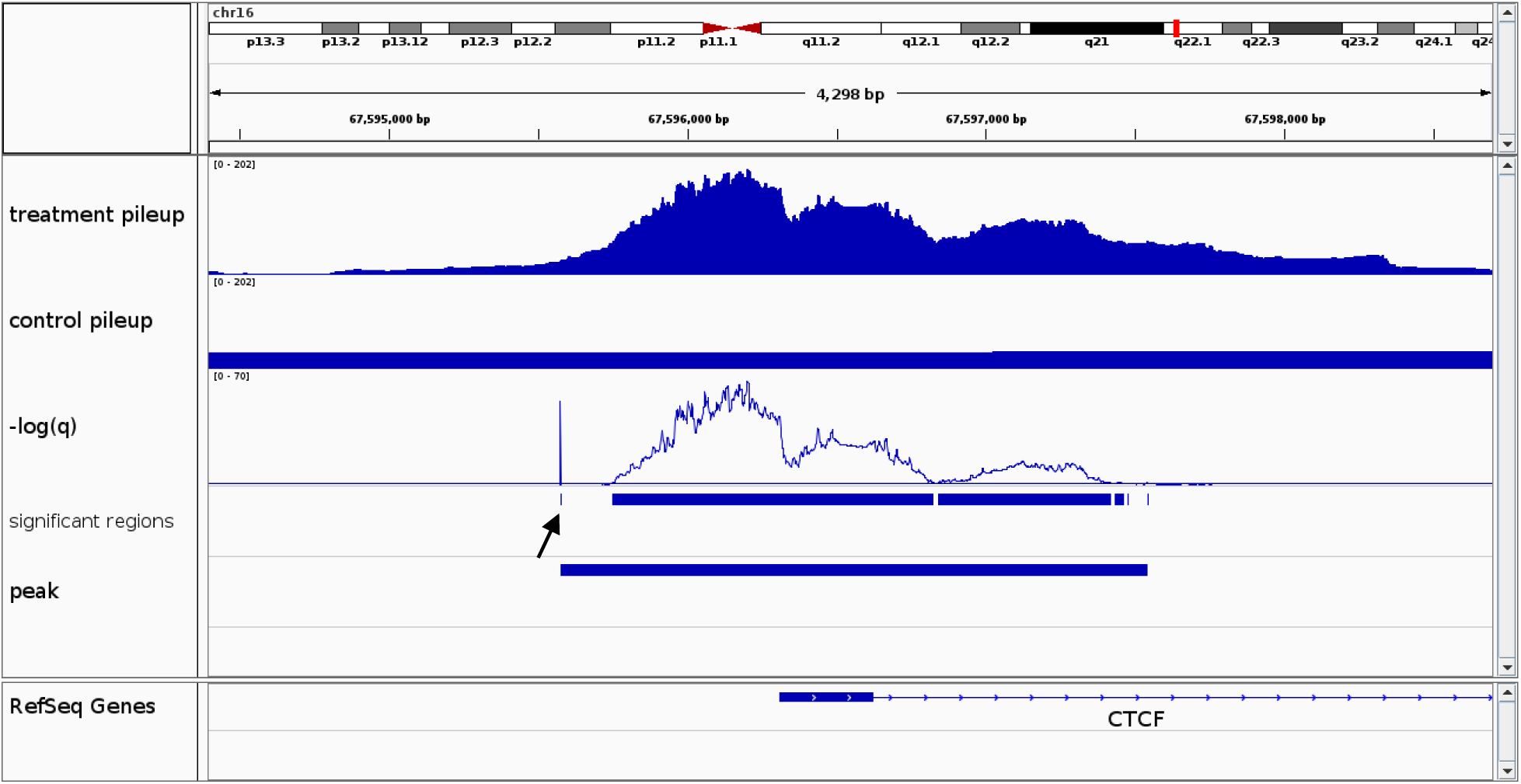
False extension of a peak due to an unresolved hash collision. For sample SRR5427886, a 1968bp peak covering the transcription start site of *CTCF* was called by MACS2. However, none of the first 175bp of the peak reached statistical significance; it was included because of a single miscalculated *q*-value (black arrow). The miscalculation was due to an unresolved hash collision in MACS2. Our modified version of MACS2 corrects this bug. Note that MACS2 was run using the dynamic lambda for the control (without the ‘--nolambda’ option).

